# MethylStar: A fast and robust pre-processing pipeline for bulk or single-cell whole-genome bisulfite sequencing data

**DOI:** 10.1101/2019.12.20.884536

**Authors:** Yadollah Shahryary, Rashmi R. Hazarika, Frank Johannes

## Abstract

**Background:** Whole-Genome Bisulfite Sequencing (WGBS) is a Next Generation Sequencing (NGS) technique for measuring DNA methylation at base resolution. Continuing drops in sequencing costs are beginning to enable high-throughput surveys of DNA methylation in large samples of individuals and/or single cells. These surveys can easily generate hundreds or even thousands of WGBS datasets in a single study. The efficient pre-processing of these large amounts of data poses major computational challenges and creates unnecessary bottlenecks for downstream analysis and biological interpretation.

**Results:** To offer an efficient analysis solution, we present MethylStar, a fast, stable and flexible pre-processing pipeline for WGBS data. MethylStar integrates well-established tools for read trimming, alignment and methylation state calling in a highly parallelized environment, manages computational resources and performs automatic error detection. MethylStar offers easy installation through a dockerized container with all preloaded dependencies and also features a user-friendly interface designed for experts/non-experts. Application of MethylStar to WGBS from human, maize and Arabidopsis shows that it outperforms existing pre-processing pipelines in terms of speed and memory requirements.

**Conclusions:** MethylStar is a fast, stable and flexible pipeline for high-throughput pre-processing of bulk or single-cell WGBS data. Its easy installation and user-friendly interface should make it a useful resource for the wider epigenomics community. MethylStar is distributed under GPL-3.0 license and source code is publicly available for download from github https://github.com/jlab-code/MethylStar. Installation through a docker image is available from http://jlabdata.org/methylstar.tar.gz

## Background

Whole-Genome Bisulfite Sequencing (WGBS) is a Next Generation Sequencing (NGS) technique for measuring DNA methylation at base resolution. As a result of continuing drops in sequencing costs, an increasing number of laboratories and international consortia (e.g. IHEC, SYSCID, BLUEPRINT, EpiDiverse, NIH ROADMAP, Arabidopsis 1001 Epigenomes, Genomes and physical Maps) are adopting WGBS as the method of choice to survey DNA methylation in large population samples or in collections of cell lines and tissue types, either in bulk or at the single-cell level [1, 2]. Such surveys can easily generate hundreds or even thousands of WGBS datasets in a single study. A broad array of software solutions for the downstream analysis of bulk and single-cell WGBS data have been developed in recent years. These include tools for data normalization such as RnBeads [3], SWAN [4], ChAMP [5], detection of differentially methylated regions (DMRs) e.g. Methylkit [6], DMRcaller [7], Methylpy [8], metilene [9], imputation of methylomes from bulk WGBS data e.g. METHimpute [10], as well as imputation of single-cell methylomes e.g. Melissa [11], deepCpG [12] and dropouts in single-cell data e.g. SCRABBLE [13].

However, these downstream analysis tools are dependent on the output of a number of data pre-processing steps, such as quality control e.g. FastQC [14], QualiMap [15], NGS QC toolkit [16], de-multiplexing of sequence reads, adapter trimming e.g Trimmomatic [17], TrimGalore [18], Cutadapt [19], alignment of reads to a reference genome and generation of methylation calls e.g. BSseeker2 [20], BSseeker3 [21], Bismark [22], BSMap [23], bwameth [24], BRAT-nova [25], BiSpark [26], WALT [27], segemehl [28]. From a computational standpoint, data pre-processing is by far the most time-consuming step in the entire bulk or single-cell WGBS analysis workflow(Fig.1). In an effort to help streamline the pre-processing of WGBS data several pipelines have been published in recent years. These include nfcore/methylseq [29], gemBS [30], Bicycle [31] and Methylpy, some of which are currently employed by several epigenetic consortia. gemBS, Bicycle and Methylpy integrate data pre-processing and analysis steps using their own custom trimming and/or alignment tools (see Table 3). By contrast, nf-core/methylseq implements well-established NGS tools, such as TrimGalore for read trimming and Bismark and bwa-meth/MethylDackel [24] for alignment. The nf-core framework is built using Nextflow [32], and aims to provide reproducible pipeline templates that can be easily adapted by both developers as well as experimentalists. Despite these efforts, the installation and execution of these pipelines is not trivial and often require substantial bioinformatic support. Moreover, managing the run times of these pipelines for large numbers of WGBS datasets (i.e. in the order of hundreds or thousands) relies on substantial manual input, such as launching of parallel jobs on a compute cluster and collecting output files from temporary folders.

**Figure 1.**
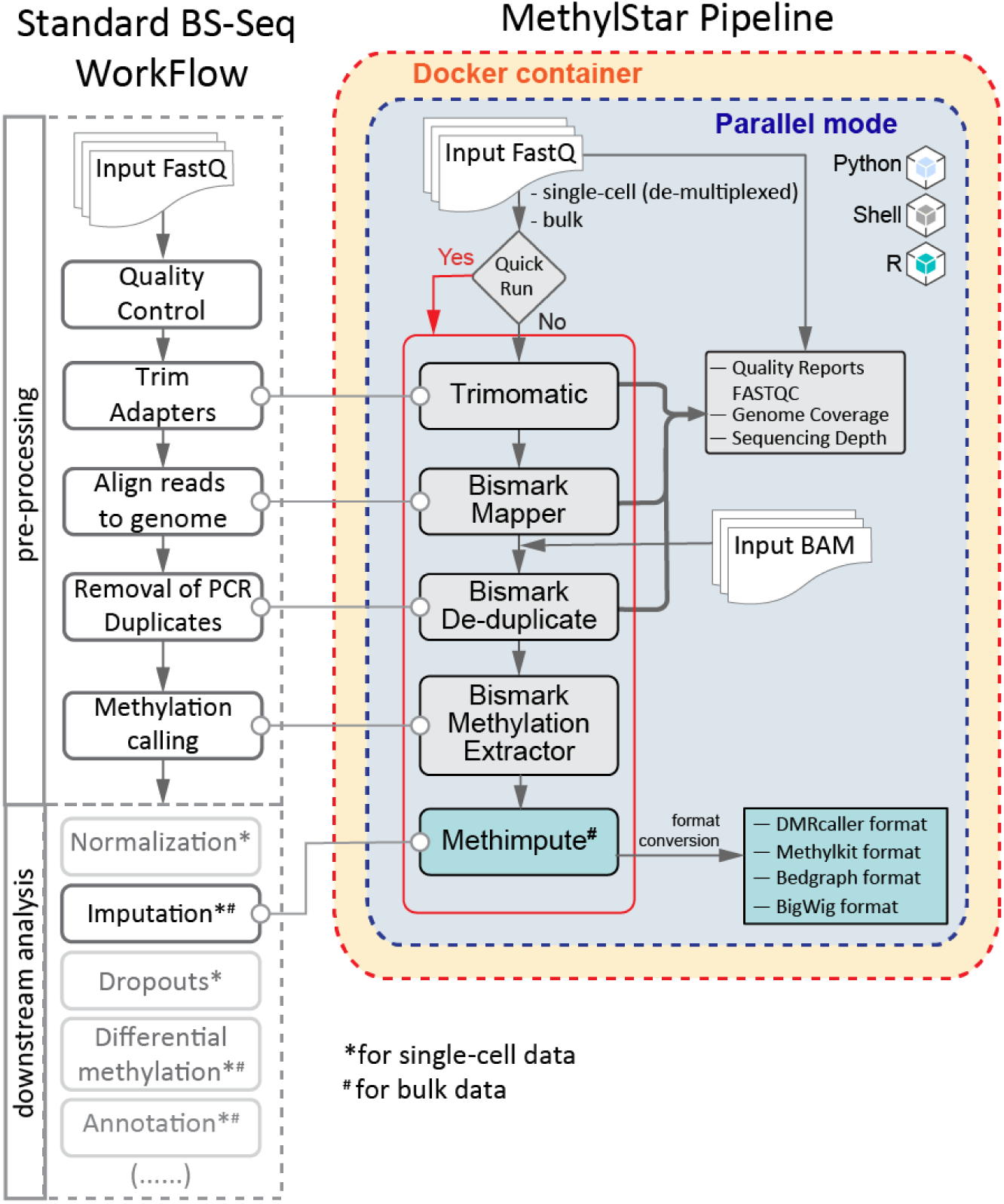
Basic workflow of MethylStar showing the pipeline architecture. The left panel shows a standard BS-Seq workflow and on the right are the different components of the MethylStar pipeline integrated as 3 different layers viz. Python, Shell and R. All steps of the pipeline have been parallelized using GNU parallel. MethylStar offers the option for “Quick run” (indicated in red) which runs all steps sequentially in one go or each component can be executed separately. MethylStar incorporates all pre-processing steps of a standard BS-Seq workflow and generates standard outputs that can be used for input into several downstream analysis tools.

In an attempt to address these issues, we have developed MethylStar, a fast, stable and flexible pre-processing pipeline for WGBS data. MethylStar integrates well-established NGS tools for read trimming, alignment and methylation state calling in a highly parallelized environment, manages computational resources and performs automatic error detection. MethylStar offers easy installation through a dockerized container with all preloaded dependencies and also features a user-friendly interface designed for experts/non-experts. Application of MethylStar to WGBS from Human, maize and Arabidopsis shows that it outperforms existing pre-processing pipelines in terms of speed and memory requirements.

## Implementation

### Core pipeline NGS components

In its current implementation, MethylStar integrates processing of raw fastq reads for both single- and paired-end data with options for adapter trimming, quality control (fastQC) and removal of PCR duplicates (Bismark software suite). Read alignment and cytosine context extraction is performed with the Bismark software suite. Alignments can be performed for WGBS and Post Bisulfite Adapter tagging (PBAT) approaches for single-cell libraries. Bismark was chosen because it features one of the most sensitive aligners, resulting in comparatively high mapping efficiency, low mapping bias and good genomic coverage [33, 34]. Finally, cytosine-level methylation calls are (optionally) obtained with METHimpute, a Hidden Markov Model for inferring the methylation status/level of individual cytosines, even in the presence of low sequencing depth and/or missing data. All the different data processing steps have been optimized for speed and performance (see below), and can run on local machines as well as on larger compute nodes.

### Pipeline architecture, optimization of parallel processes and memory usage

The pipeline architecture comprises three main layers (Fig.1). The first layer is the interactive command-line user interface implemented in Python to simplify the process of configuring software settings and running MethylStar. Easy navigation through this interface allows non-experts to run large batches of samples without having to type commands at the terminal. The second layer consists of shell scripts, which handle low-level processes, efficiently coordinates the major software components and manages computational resources. The final layer is implemented in R, and is used to call METHimpute and to generate output files that are compatible with a number of publicly available DMR-callers such as Methylkit, DMRcaller and bigWig files for visualization in Genome Browsers such as JBrowse [35]. All outputs are provided in standard data formats for downstream analysis.

All components/steps of the pipeline including adapter trimming, read alignment, removal of PCR duplicates and generation of cytosine calls have been parallelized using GNU Parallel [36] (Fig.1). The user can either set the number of parallel jobs manually for each pipeline component, or can opt to use the inbuilt parallel option. The inbuilt parallel implementation is available under the “Quick Run” option, which detects the number of parallel processes/jobs automatically for each pipeline component based on available system cores/threads and memory, thus allowing the user to run the entire steps of the pipeline in one go. In the parallel implementation of the Bismark alignment step, we include the genome size (in base pairs) as an additional factor while optimizing computational resources. For example, while running paired-end reads from *A. thaliana* with a genome size of ~135 Mb on a system with 88 cores and 386 GB RAM we optimally set the number of jobs to 4. This setting allocates (4 jobs × 8 files/threads) =32 threads to Bowtie2 and (4 jobs × 8 files/threads × 2) =64 threads to the bismark alignment tool (default no. of threads fixed to 8 in the internal bismark parallel argument). In this way, the maximum number of threads never exceeds the total number of available cores, which in turn allows other jobs such as file compression, I/O operations to be performed simultaneously.

Under the “Quick Run” option we have parallelized R processes such as the extraction of methylation calls from BAM files (post PCR duplicates removal) by bypassing the Bismark methylation extractor step and by passing these calls directly onto METHimpute for imputation of missing cytosines (Fig.1). In the parallelization of R processes we allocate even fewer number of threads (=3 threads in our system with 88 cores and 386 GB RAM), as these processes (in our case extracting and sorting bam files) are resource hungry and tend to load all its objects into memory. This allows for faster processing times and efficient management of resources without crashing the entire parallel process. In addition, we have introduced checkpoints for each individual component of the pipeline so that a job can be resumed easily in the unlikely case of system failure or any kind of user interruption.

### Running MethylStar

The user can choose to run each pipeline component individually, and customize software settings at each step by editing the configuration file which is available as an option through the interactive command-line user interface. The user interface displays the available options as a list, and users can execute specific pipeline steps by simply typing the index of their choice. Some of the key configuration parameters include setting file paths to input and output data, as well as options for handling large batches of samples, conversions to required file formats and deletion of auxiliary files that were generated during intermediate analysis steps. Our interactive user interface aids in the fast execution of complex commands and will be particularly effective for users who are less familiar with command line scripting. As an alternative, MethylStar also features a “Quick Run option”, which allows the user to run all pipeline steps in one go using default configuration settings (Fig.1).

### Installation and documentation

MethylStar can be easily installed via a Docker image. This includes all the softwares, libraries and packages within the container, and thus solves any dependency issues. Advanced users can edit the existing docker container and build their own image.

Detailed description about installation and running the pipeline is available at https://github.com/jlab-code/MethylStar

## Results and Discussion

### Benchmarking of speed

To demonstrate MethylStar’s performance we analyzed bulk WGBS data from a selection of 200 *A. thaliana* ecotypes (paired-end, 295GB, ~8.63× depth, 85.66% genome coverage, GSE54292), 75 maize strains (paired-end, 209GB, ~0.36× depth, ~22.12% genome coverage, GSE39232) and 88 Human H1 cell lines (single-end, 82GB, ~0.12× depth, ~10.62% genome coverage, GSM429321). MethylStar was compared with Methylpy, nf-core/methylseq and gemBS. All pipelines were run with default parameters on a computing cluster with a total of 88 cores (CPU 2.2 GHz with 378 GB RAM). Speed performance was assessed for a series of batch sizes (*A. thaliana*: 50, 100, 150, 200 samples; human H1 cell line: 22, 44, 66, 88 samples; maize: 15, 30, 45, 60, 75 samples) and was restricted to a fixed number of jobs (=32), see Fig. 2A-C. Although gemBS achieved the fastest processing times for the *A. thaliana* samples, MethylStar clearly outperformed the other pipelines when applied to the more complex genomes of maize and human, which are computationally more expansive and resource-demanding (Fig. 2B-C). For instance, for 88 human WGBS samples (82GB of data), MethylStar showed a 75.61% reduction in processing time relative to gemBS, the second fastest pipeline (909 mins vs. 3727 mins). Extrapolating from these numbers, we expect that for 1000 human WGBS samples, MethylStar could save about ~22.24 days of run time (4× faster). To show that MethylStar can also be applied to single-cell WGBS data, we analyzed DNA methylation of 200 single cells from human early embryo tissue (paired-end, 845GB, ~0.38× depth, ~9.97% genome coverage, GSE81233) split into batches of 100 and 200, see Fig. 2D. MethylStar’s processing times increased linearly with batch size (i.e. number of cells). For 200 cells, MethylStar required only 4227 mins, thus making it an efficient analysis solution for deep single-cell WGBS experiments.

**Figure 2.**
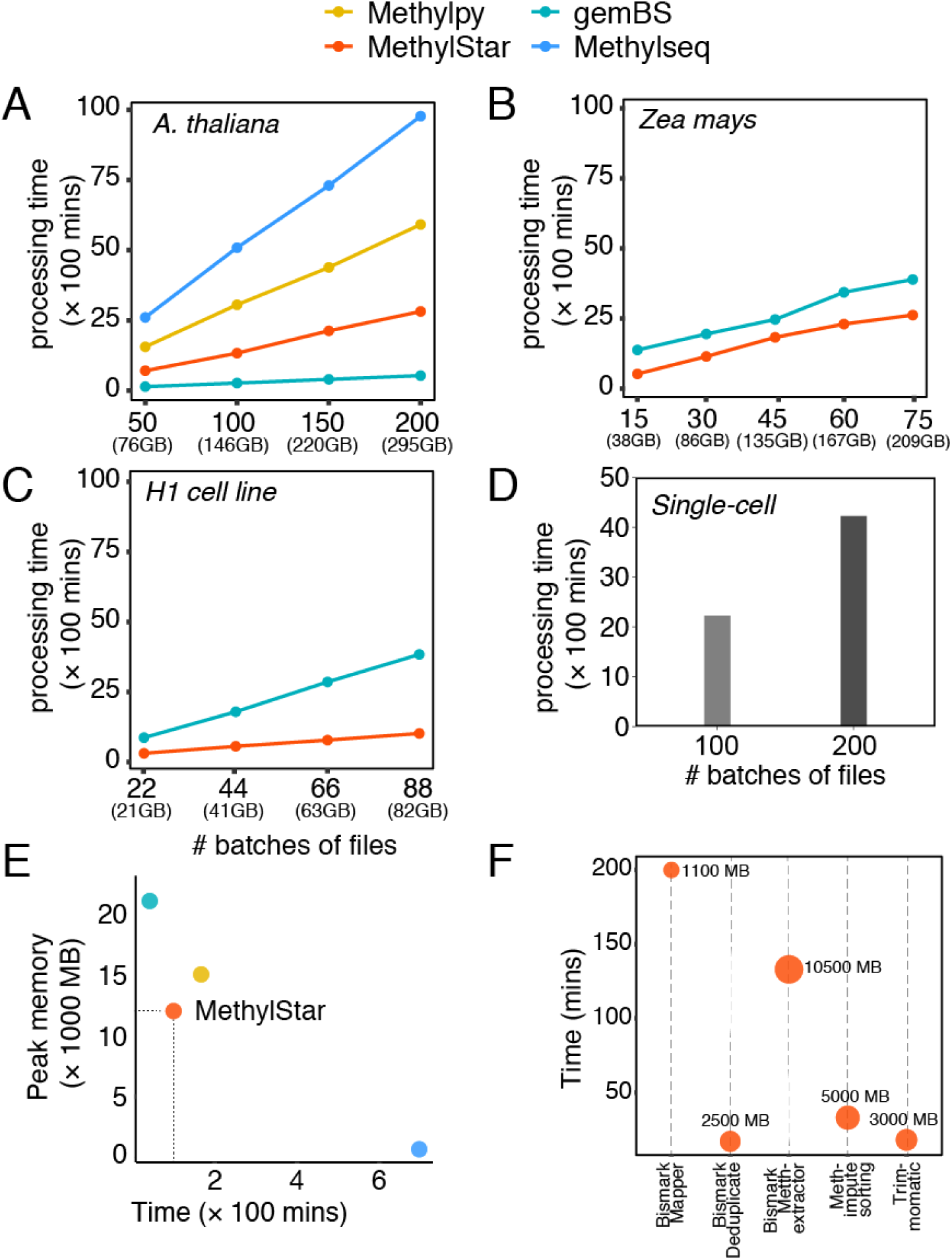
Performance of MethylStar as compared with other BS-Seq analysis pipelines viz. Methylpy, nf-core/methylseq and gemBS in **(A)** A. thaliana **(B)** maize **(C)** H1 cell line and **(D)** scBS-Seq samples. CPU processing time taken by METHimpute was not included in the current benchmarking process as there is no equivalent method in the other pipelines to compare with. Because of the very long run times observed for the A. thaliana data, Methylpy and Methylseq were no longer considered for benchmarking of speed in maize and H1 cell line samples. All pipelines were run using 32 jobs. **(E)** Peak memory usage as a function of time for 10 random A. thaliana samples. **(F)** Time taken by each component of MethylStar. X-axis shows the individual components of MethylStar and on the y-axis is the time in mins. The size of the dot indicates the peak memory usage by each component.

**Figure 3.**
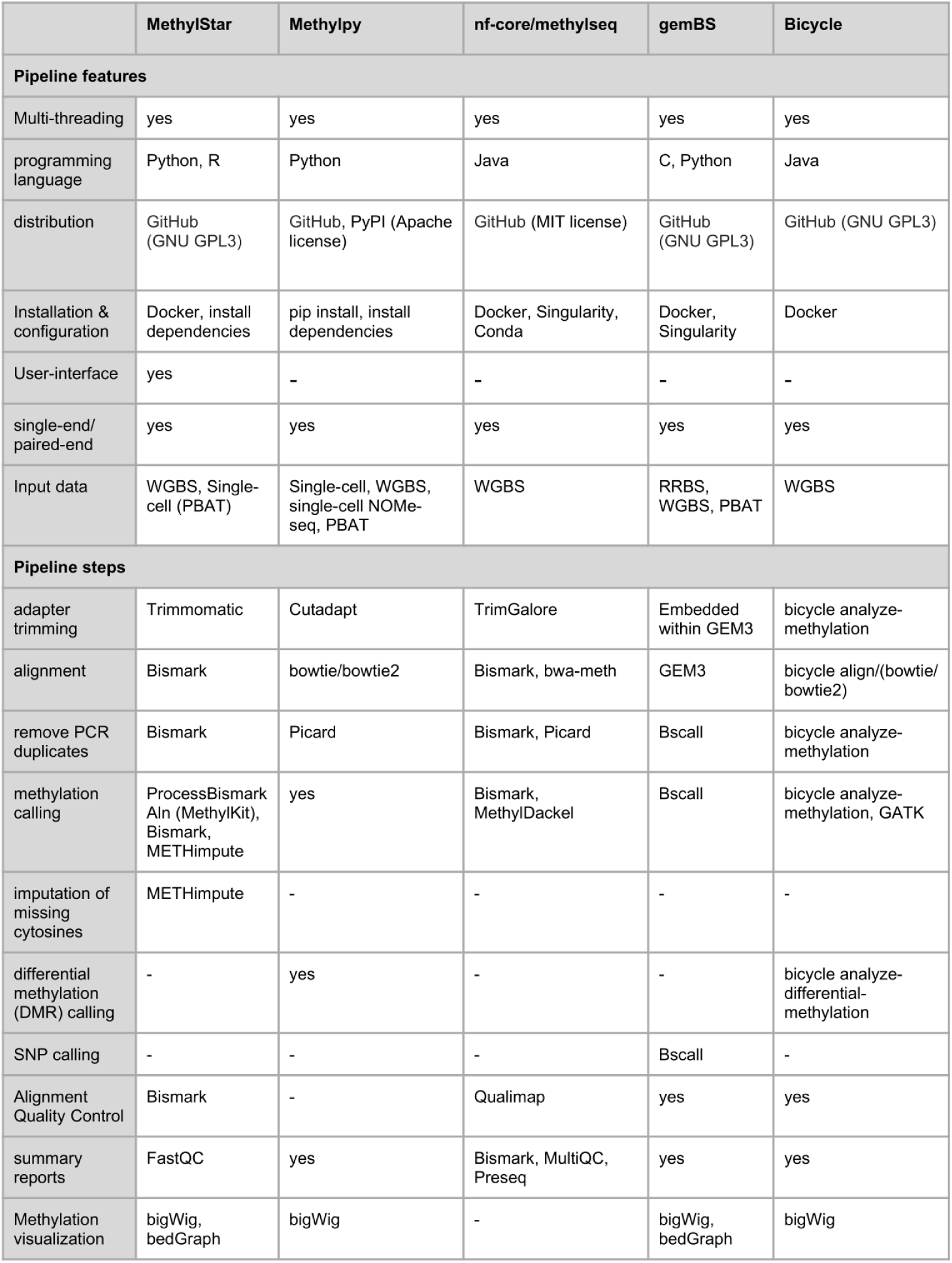
Table showing different features of MethylStar as compared to other BS-seq pipelines

### Memory usage statistics

Along with benchmarking of speed, we also evaluated the performance of the MethylStar, gemBS, nf-core/methylseq and Methylpy pipelines in terms of system memory utilization using the MemoryProfiler [37] python module (Fig. 2E). We assessed the CPU time versus peak/max memory of all the 4 pipelines (default settings) on a computing cluster (specifications above). For 10 random samples from the above *A. thaliana* benchmarking dataset (paired-end, 16GB, GSE54292) MethylStar and Methylpy showed the best balance between peak memory usage (~12000 MB and ~15000 MB, respectively) and total run time (~100 mins and 167 mins, respectively). In contast, nf-core/Methylseq and GemBS exhibited strong trade-offs between memory usage and speed, with nf-core/Methylseq showing the lowest peak memory usage (~700 MB) but the longest CPU time (~697 mins), and GemBS the highest peak memory usage (~21000 MB) but the shortest run time (~42 mins) (Fig. 2E). Furthermore, we inspected the time taken by each individual component of MethylStar. Bismark alignment was the most time consuming step of the pipeline but required the lowest peak memory usage (~1100MB) of all the steps, indicating that our parallel implementation of the Bismark alignment step can be very effective in handling large numbers of read alignments with low memory requirements (Fig. 2F). We further benchmarked memory usage using 10 random samples from the above maize dataset (paired-end, 23GB, GSE39232). For this analysis, we focused on gemBS and MethylStar due to their shorter processing times for these datasets as compared to nf-core/Methylseq and Methylpy. For these maize dataset, gemBS’s peak memory usage was ~110000 MB as compared to ~81000 MB for MethylStar (~1.3 times less memory) with a total run time of 667 mins and 421 mins, respectively. Taken together, these benchmarking results clearly show that MethylStar exhibits favorable performance in terms of processing time and memory, and that it is therefore an efficient solution for the pre-processing of large numbers of samples even on a computing cluster with limited resources.

## Conclusion

MethylStar is a fast, stable and flexible pipeline for the high-throughput analysis of bulk or single-cell WGBS data. Its easy installation and user-friendly interface should make it a useful resource for the wider epigenomics community.

